# Deregulated mito-nuclear communication alters chromatin plasticity and differentiation potential of mesenchymal stem cells upon ageing

**DOI:** 10.1101/2020.04.02.022293

**Authors:** Andromachi Pouikli, Swati Parekh, Monika Maleszewska, Maarouf Baghdadi, Ignacio Tripodi, Chrysa Nikopoulou, Kat Folz-Donahue, Yvonne Hinze, Andrea Mesaros, Patrick Giavalisco, Robin Dowell, Linda Partridge, Peter Tessarz

## Abstract

Ageing is accompanied by a general decline in the function of many cellular pathways, with metabolic alterations, epigenetic modifications, and stem cell exhaustion representing three important hallmarks of the ageing process. However, whether these pathways are causally or functionally related at a molecular level remains poorly understood. Here, we use bone marrow-derived mesenchymal stem cells (MSCs) isolated from young and old mice to address how age-dependent changes in metabolism and epigenetics are linked and how they impact on the ageing transcriptome and differentiation potential. Given that MSCs maintain specific age-associated properties even under prolonged culture conditions, such as the age-dependent decrease in osteogenic differentiation, they are an excellent model to investigate *in vitro* the connection of ageing hallmarks on a mechanistic level. In this study, we demonstrate that upon ageing, osteogenic potential of MSCs declines as a consequence of deregulated mito-nuclear communication, mediated by decreased levels of the citrate carrier (CiC). Age-dependent down-regulation of CiC results in acetyl-CoA trapping within mitochondria, hypo-acetylation of histones and chromatin compaction. Together, these changes lead to an altered transcriptional output and are responsible for the reduced differentiation capacity into osteoblasts. Strikingly, short-term supplementation of aged cells with acetate, an exogenous source for cytosolic acetyl-CoA production, rescues not only the age-associated reduction of histone acetylation, but also the osteogenesis defect, representing a potential target for *in vitro* MSC rejuvenation.

## INTRODUCTION

Stem cell exhaustion, a result of reduced self-renewal capacity and imbalanced differentiation potential, is a well-established hallmark of the ageing process (López-Otín et al. 2013). Bone marrow mesenchymal stem cells (MSCs) have been shown to play an important role upon ageing due to their ability to regenerate bone by giving rise to adipocytes, chondrocytes and osteoblasts (Dominici et al. 2006; da Silva Meirelles, Caplan, and Nardi 2008; Caplan 2008). Aged MSCs show decreased capacity to differentiate into the osteogenic and chondrogenic lineages. This feature has been linked to increased fat content in the bone marrow upon ageing and concomitantly higher risk of osteoporosis and fractures (Kim et al. 2012; S. Zhou et al. 2008). This change in fate commitment is also known to occur in other stem cell populations upon ageing, such as hematopoietic stem cells, which show a biased differentiation potential towards the myeloid lineage (Akunuru and Geiger 2016).

Chromatin architecture influences stem cell fate decisions and cell differentiation is often accompanied by chromatin re-arrangements (Dixon et al. 2015). Research over the last few years has revealed that ageing manifests itself - among others - by changes in chromatin architecture and subsequently by alterations in gene expression across cell types and tissues (Benayoun, Pollina, and Brunet 2015; Booth and Brunet 2016). Thus, epigenetic changes are considered an important hallmark of the ageing process (López-Otín et al. 2013). Recently, partial reprogramming in various tissues was found to result in tissue regeneration and to prolong lifespan in premature ageing models (Ocampo et al. 2016), strongly suggesting that interfering with epigenetic mechanisms provides a way to intervene in the ageing process.

Metabolism has also been strongly implicated to contribute to the maintenance of stem cell states (Ito and Suda 2014). In this context, it is important to highlight that metabolism and chromatin are heavily intertwined (Kaelin and McKnight 2013; Lu and Thompson 2012; Reid, Dai, and Locasale 2017; Etchegaray and Mostoslavsky 2016). More precisely, intracellular metabolism provides metabolites that serve as essential cofactors and substrates for chromatin-modifying enzymes and their availability can strongly impact the activity of these enzymes. Many of these metabolites are generated within mitochondria and this establishes a tight mitochondrial-nuclear connection (Quirós, Mottis, and Auwerx 2016). For instance, during histone acetylation, histone acetyltransferases (HATs) transfer acetyl-groups from acetyl-CoA to lysine residues of histones. In mammals, most of the cytosolic acetyl-CoA is derived from the mitochondria-produced citrate, via a reaction catalysed by the enzyme ATP-citrate lyase (ACLY) (Srere 1959).

Like many adult stem cell populations, MSCs reside in niches characterised by low oxygen concentration (Ito and Suda 2014). In particular, the bone marrow niche was found to contain 1.3-4.2% oxygen (Spencer et al. 2014). Thus, MSCs rely on anaerobic glycolysis for their energy production. However, glycolysis is not only the preferred metabolic pathway to produce energy, but it has been also suggested to maintain MSCs in a multipotent state (Suda, Takubo, and Semenza 2011). On the contrary, increased mitochondrial respiration and oxidative phosphorylation (OXPHOS) have been demonstrated to occur during differentiation of MSCs and to facilitate lineage-specific commitment (Shyh-Chang, Daley, and Cantley 2013).

Interestingly, MSCs purified from old individuals maintain specific, age-related properties, such as the decreased differentiation potential into the osteogenic lineage, even after prolonged culture *in vitro*, suggesting stable inheritance of epigenetic states. Thus, MSCs represent an ideal model to systematically study *in vitro* the age-related alterations in chromatin, metabolism and cell fate on a mechanistic level.

In this study we demonstrate that osteogenesis defects observed in aged MSC originate from age-associated changes in the chromatin landscape. We show that upon ageing histone acetylation states are strongly impacted by an altered metabolic profile and we identify impaired acetyl-CoA export from mitochondria to the cytoplasm due to lower levels of citrate carrier as a critical component of this process. Impressively, circumventing defective acetyl-CoA export from mitochondria by supplementing medium of aged cells with acetate rescues loss of histone acetylation and the impaired osteogenic differentiation of aged MSCs. Collectively, our results establish a tight, age-dependent connection between metabolism, chromatin and stemness and identify citrate carrier as a critical mediator of mitochondrial-nuclear communication during ageing of MSCs.

## RESULTS

### Aged MSCs exhibit low proliferation and differentiation capacity

We isolated MSCs from the endosteum of young (∼ 3-5 months old) and old (∼ 18-22 months old) mice, following a published protocol outlined in Figure 1A (Houlihan et al. 2012). This strategy purifies a homogenous population of bone-marrow MSCs expressing high levels of Sca-1 and CD140a mesenchymal stem cell markers (Figure S1A, from here on referred to as MSCs). MSCs isolated using this protocol exhibit enhanced growth potential and robust tri-lineage differentiation capacity, compared to standard plastic adherence-selected MSCs. In addition, the selective isolation of MSCs based on specific surface markers avoids cellular contamination that could potentially interfere with downstream analysis (Houlihan et al. 2012). Furthermore, these MSCs have been shown to maintain their tri-lineage potential after transplantation, suggesting that they represent an excellent model to study MSC biology (Morikawa et al. 2009). Since only 500-1000 MSCs could be isolated per mouse following this procedure, we modified this protocol by adding the possibility for outgrowth of more MSCs from bones *in vitro*, before cell sorting, in order to acquire sufficient cells for biochemical assays (∼40,000 cells per mouse). Taking into consideration the variability between individual mice, we pooled together cells obtained from three different animals, in each biological replicate. Purified MSCs were characterized by the expression of mesenchymal stem cell surface markers, while being devoid of hematopoietic markers (Figure S1A).

**Figure 1.**
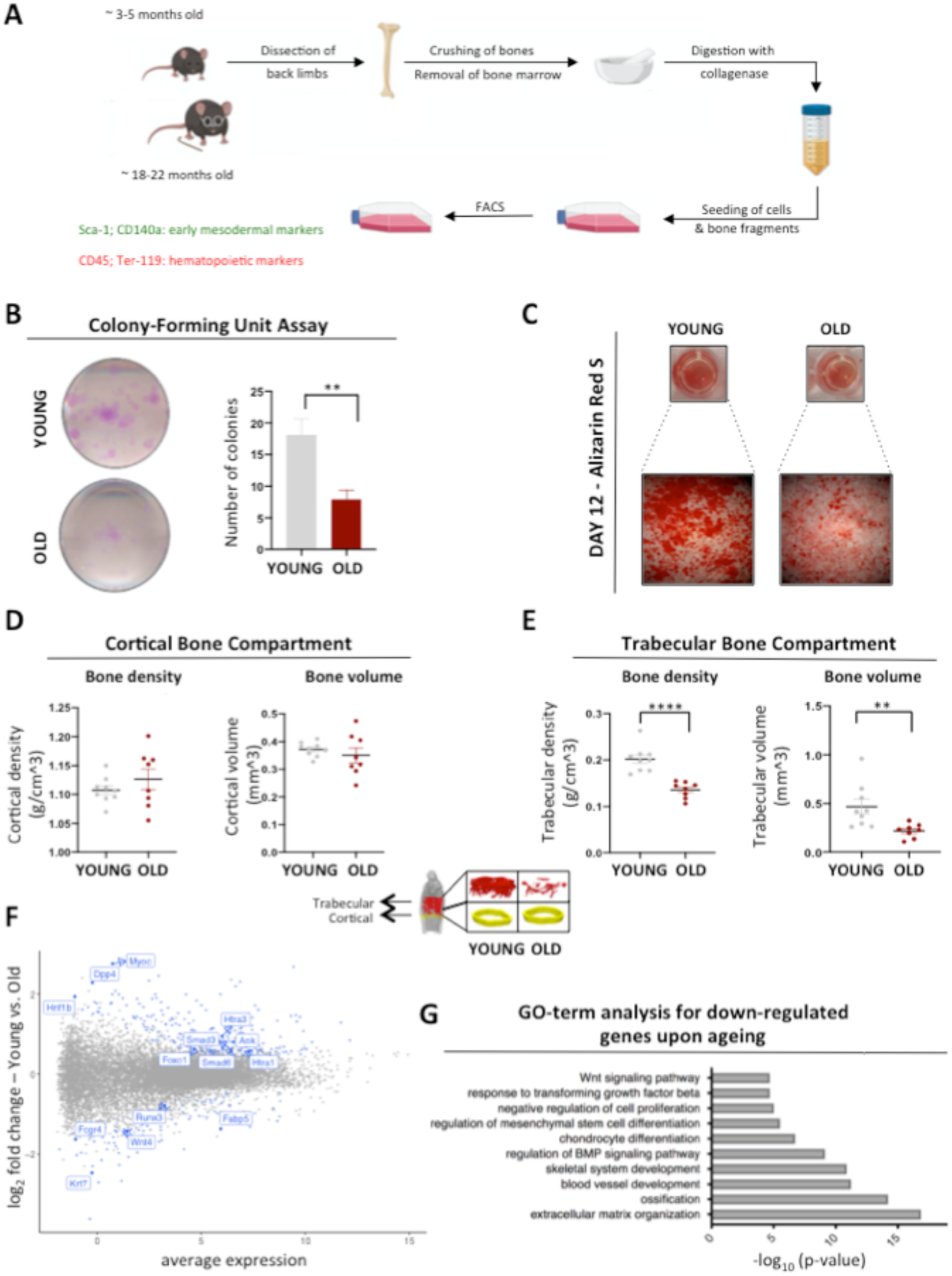
MSCs lose their proliferation and differentiation potential upon ageing. **(A)** Schematic representation demonstrating the isolation protocol of bone marrow Mesenchymal Stem Cells (MSCs) from the back limbs of young (∼3-5 months old) and old (∼18-22 months old) mice. After collection of the limbs, clean bones were cut into small pieces, which were then treated with collagenase for 1h at 37°C. Cells and bone fragments were seeded on flasks and incubated for 10 days under 2% O2. On day 10, we performed a cell sorting analysis using flow cytometry; we sorted for the CD45-/Ter-119-double negative mesenchymal stem cells. **(B)** Representative images and quantification of Colony Forming Unit assay for young and aged MSCs, after culturing them for 10-14 days; (n=3 biological replicates for each age group). **(C)** Representative images of Alizarin-Red-S staining performed on young and aged MSCs, 12 days after ostegenesis induction; (n=3 biological replicates for each age group). Images were acquired using bright-field microscopy. **(D and E)** Measurement of bone density and bone volume of the cortical and trabecular bone compartments from the femurs of young and old animals; (n=9 biological replicates for young mice; n=8 biological replicates for old mice). **(F and G)** MA-plot and GO-term analysis for down-regulated genes upon ageing, identified by RNA-seq. Data are presented as mean ± SEM. Statistical significance was determined using student’s t-Test; **p-value < 0,01; ****p-value <0,0001. See also Figure S1.

We confirmed here that these cells showed three-lineage differentiation capacity, being able to give rise to adipocytes, chondrocytes and osteoblasts (Figure S1B-D), as has been previously reported for cells obtained using this protocol (Houlihan et al. 2012). This clearly demonstrates that our purification strategy enables successful isolation of multipotent MSCs. These cells were cultured exclusively under 2% oxygen, similar to the oxygen concentration found in their niche (Spencer et al. 2014).

One characteristic described for plastic-adherent MSCs is the loss of stemness when isolated from older individuals. To test whether our homogenous MSC’s population shows the same phenotype, we performed proliferation and differentiation assays. Indeed, MSCs isolated from young mice produced more and bigger colonies in comparison to MSCs from old mice (Figure 1B), indicating that the self-renewal ability was reduced upon stem cell ageing.

Next, we sought to investigate changes in the differentiation capacity of young and aged cells. While the adipogenic differentiation capacity was only mildly affected during ageing (Figure S1E), the ability of aged MSCs to differentiate into osteocytes was significantly decreased (Figure 1C). These observations prompted us to correlate our *in vitro* findings with potential *in vivo* changes in the bone quality upon ageing. Hence, we measured bone mineral density and bone volume in cortical and trabecular bone compartments of femurs from young and old mice. Interestingly, we observed decreased trabecular density and volume in old mice, whereas cortical density and volume remained unaffected during ageing (Figures 1D and 1E). To understand whether the observed changes in bone quality and stem cell potency were due to ageing-driven alterations in the transcriptional output, we performed RNA-sequencing (Figures 1F and 1G).

Interestingly, we found that Wnt4 and Runx3 genes were among the top down-regulated genes in aged cells; both genes have been reported to contribute significantly to stem cell potency because of their role in cell proliferation and osteogenesis, respectively (Saito et al. 2015; Chang et al. 2007). On the other hand, Dpp4 was one of the most up-regulated genes in aged MSCs. This gene encodes a protease that has been recently shown to prevent bone regeneration, whereas inhibition of Dpp4 accelerates tibia fracture healing and increases the abundance of osteogenic progenitors (Ambrosi et al. 2017). Together, these results indicate that the age-related differentiation bias against osteogenic and chondrogenic lineages is fully supported by alterations in the transcriptional profile of aged MSCs.

### Chromatin re-arrangements in MSCs upon ageing

Chromatin architecture plays a critical role in the regulation of gene transcription(B. Li, Carey, and Workman 2007). Therefore, we investigated whether age-associated alterations in the chromatin landscape were responsible for the transcriptional changes presented above. We measured global chromatin accessibility by Assay for Transposase-Accessible Chromatin using sequencing (ATAC-seq, Figure S2) (Buenrostro et al. 2013) on young and aged MSCs.

This assay monitors chromatin accessibility on a genome-wide scale. We found that upon ageing 14750 sites in the genome changed significantly in accessibility status (FDR < 0.05); in particular, ∼6500 sites became more accessible, while ∼8250 sites became less accessible (Figure 2A). Surprisingly, we observed that global chromatin accessibility decreased significantly with age (Figure 2B). To further analyze differences in the chromatin structure, we plotted accessibility over the transcription start site (TSS) as a metaplot (Figure 2C). We observed that gene promoters in aged cells exhibited a strong decrease in chromatin accessibility. In order to understand if the observed chromatin changes could explain the changes in transcriptional output, we integrated the outcome of ATAC-seq and RNA-seq analysis using Metascape. Metascape allows for a global overview of the transcriptional activity by identifying common gene ontology (GO) enrichment, between ATAC-seq and RNA-seq datasets (Y. Zhou et al. 2019). Common GO-terms converged on cell proliferation and osteogenesis / bone regeneration processes (Figure 2D). Importantly, our data also suggested de-regulation of Wnt and BMP signaling, two pathways of critical importance for bone development (Deschaseaux, Sensébé, and Heymann 2009; Stewart et al. 2014).

**Figure 2.**
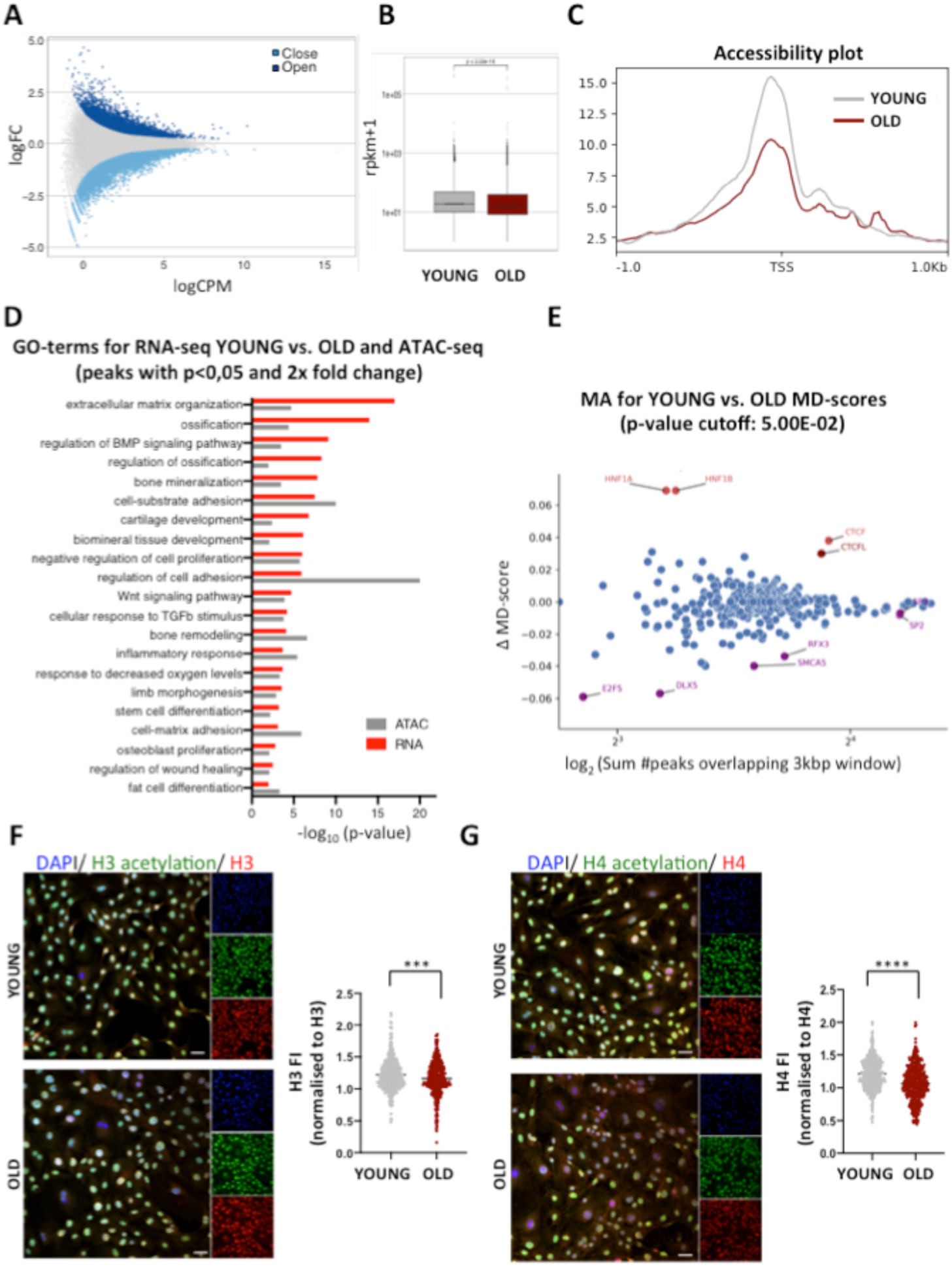
Chromatin compaction and histone hypo-acetylation upon ageing. **(A)** MA-plot showing opening and closing peaks with age as determined by ATAC-seq. **(B)** Overall genome accessibility measured by ATAC-seq. **(C)** Metaplot of ATAC-seq reads over the transcription start site (TSS) of all genes. **(D)** Common GO-terms that overlap in ATAC- and RNA-seq datasets were determined using the Metascape algorithm. **(E)** DAStk analysis of the ATAC-seq data to predict transcription factor activity. **(F and G)** Representative images and quantification after immunofluorescence staining of young and aged cells, using antibodies against histone H3 acetylation and H4 acetylation. Levels of Histone H3 and Histone H4 fluorescence intensity were used as an internal control, respectively, for normalization. Nuclei were stained with DAPI. Scale bars, 75μm. Data are presented as mean ± SEM. Statistical significance was determined using student’s t-Test; ***p-value < 0,001; ****p-value <0,0001. See also Figure S2.

Finally, we used our ATAC-seq data to interrogate if specific transcription factors were responsible for the observed chromatin re-arrangements. To answer this question, we applied the Differential ATAC-seq toolkit (DAStk) (Tripodi, Allen, and Dowell 2018) on our ATAC-seq dataset. This approach uses ATAC-seq datasets to predict transcription factor activity. DAStk analysis identified several transcription factors and chromatin remodelers that could have contributed to the accessibility changes observed by ATAC-seq (Figure 2E).

The most significant hit was CTCF, which was predicted to be more active in the aged cells. By contrast, higher activity of the Dlx5 and E2F5 transcription factors was implicated in young cells. These transcription factors are involved in cell proliferation and osteogenesis, respectively, and correlate with the observed age-related effects on proliferation ability and osteogenic differentiation capacity (Figures 1B and 1C). In addition to transcription factors, modifications on DNA and histones shape chromatin states. In particular, histone acetylation neutralizes the positive charge of the lysine side-chains of histones, weakening the contact between histones and DNA (Tessarz and Kouzarides 2014). We thus investigated if alterations in histone acetylation profiles also contributed to changes in chromatin accessibility. In line with the overall age-associated chromatin compaction, we observed significantly decreased levels of histone H3 and histone H4 acetylation in MSCs from old mice (Figures 2F and 2G). Together, our ATAC-seq data and immunofluorescence results suggest that ageing is accompanied by histone hypo-acetylation, concomitantly with a decrease in chromatin accessibility and transcriptional changes that are responsible for the lower proliferation and differentiation capacity of aged MSCs.

### Aged MSCs switch from glycolysis to fatty acid oxidation as an energy source

Our findings regarding age-associated decrease in the levels of histone acetylation (Figures 2F and 2G) prompted us to investigate how histone acetylation profile was established in young and aged MSCs and whether other cellular processes were involved in this phenotype. In particular, we focused on metabolism, because metabolic pathways can directly impact DNA and histone modifications (Kaelin and McKnight 2013; Lu and Thompson 2012; Reid, Dai, and Locasale 2017; Etchegaray and Mostoslavsky 2016). For instance, local availability of acetyl-CoA directly influences levels of histone acetylation (Wellen et al. 2009; Cai et al. 2011).

Therefore, we measured acetyl-CoA concentration in young and aged MSCs and observed a strong increase in acetyl-CoA levels in aged MSCs (Figure 3A). This result was unexpected and in contradiction to the histone hypo-acetylation observed in MSCs from old mice. Thus, we were motivated to investigate the source of the acetyl-CoA increase. Given that in MSCs anaerobic glycolysis is the main energy source and that a direct connection between histone acetylation and glycolytic rate has been previously described (Cluntun et al. 2015), we first investigated ageing-driven changes in glycolysis. We measured glucose uptake and lactate secretion as a proxy for the glycolytic flux (TeSlaa and Teitell 2014). We found that MSCs derived from young mice consumed glucose and concomitantly secreted lactate (Figures 3B and 3C). However, MSCs purified from old mice displayed lower glucose uptake and subsequently lower lactate secretion and higher pH in their media (Figures 3B and 3C; Figure S3A), indicating that MSCs from old mice performed glycolysis to a lower extent than the young MSCs. Due to changes in the metabolic profile of aged MSCs, we next sought to further metabolically characterize MSCs upon ageing and we focused on mitochondrial metabolism; we monitored changes in the mitochondrial networks by observing cells under the electron microscope and also after staining them with TOMM20. Both experimental approaches revealed that aged cells contained fragmented mitochondria, which formed less complex and less tubular networks (Figure S3B), a phenotype which has been described for several cell types upon ageing (Sharma et al. 2019). However, we found that although ageing affects mitochondrial structure, it doesn’t have an impact on mitochondrial DNA content and mitochondrial mass (Figures S3C and S3D). To test whether the age-associated alterations in mitochondrial morphology are accompanied by changes in mitochondrial function we performed targeted metabolomics. Interestingly, we observed that many TCA cycle intermediates and TCA cycle-related amino acids were slightly enriched in aged cells (Figure 3D). This establishes a trend towards more TCA cycle intermediates in aged cells, suggesting an ageing-driven rewiring of the Krebs cycle which results in higher production and/or lower consumption of the intermediate metabolites.

**Figure 3.**
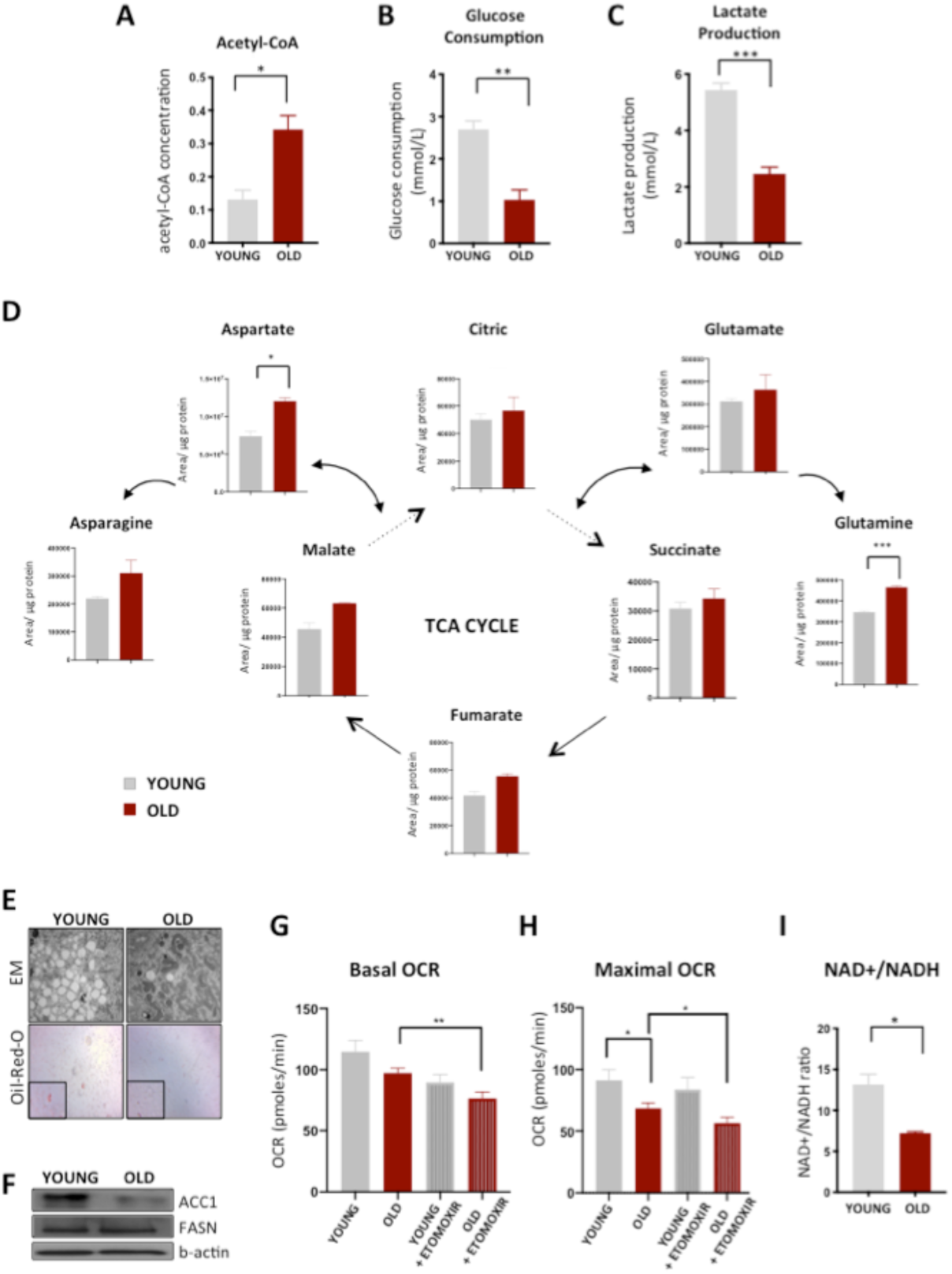
Aged MSCs down-regulate glycolysis and up-regulate β-oxidation of fatty acids. **(A)** Measurement of acetyl-CoA levels in young and aged MSCs using Mass Spectrometry analysis; (n=3 biological replicates for each age group). **(B and C)** Glucose consumption and Lactate production measured in the a-MEM media of young and aged cells using the Vi-Cell MetaFLEX instrument; (n=3 biological replicates for each age group). **(D)** Metabolomic analysis of TCA-cycle intermediates and TCA-cycle related amino acids by Mass Spectrometry; (n=3 biological replicates for young mice; n=2 biological replicates for old mice). **(E)** Representative images of lipid droplets in young and aged cells. Images were acquired by electron microscopy (top) and by bright-field microscopy, after staining of the neutral lipids with Oil-Red-O (bottom); (n=3 biological replicates for each age group). **(F)** Representative immunoblots for ACC1 and FASN proteins. β-actin was used as a loading control (n=2 for each age group). **(G and H)** Basal and Maximal Oxygen Consumption Rates (OCR) in young and aged MSCs, treated with or without 100μM Etomoxir for 1hr, prior to the assay. Measurements were made using the SeaHorse XF24 extracellular flux analyzer instrument; (n=3 biological replicates for each age group). **(I)** Metabolomic analysis of NAD+/NADH levels using Mass Spectrometry and graph of their ratio (n=3 biological replicates for each age group). Data are presented as mean ± SEM. Statistical significance was determined using student’s t-Test; *p-value < 0,05; **p-value < 0,01; *** p-value <0,001. See also Figure S3.

Finally, we investigated lipid metabolism and fatty acid oxidation (FAO) as a potential source for the higher acetyl-CoA levels upon ageing. Surprisingly, both by electron microscopy and after Oil-Red-O staining of neutral lipids, we observed that aged cells contained fewer lipid droplets than young cells (Figure 3E). Lipid droplets are the intracellular storage sites of neutral lipids (Walther and Farese 2012; Brasaemle 2007; Bickel, Tansey, and Welte 2009) and tend to accumulate in cells that do not use fatty acids to produce energy. The finding that aged cells contained fewer lipid droplets could be explained either by lower fatty acid biosynthesis or by increased lipid droplet consumption and thus higher FAO. Interestingly, although FASN protein levels did not change upon ageing, protein levels of the ACC1 enzyme, which is important for the initiation of the fatty acid biosynthesis program, were strongly decreased in aged cells (Figure 3F). These results pointed towards impaired lipogenesis upon ageing. In parallel, we measured oxygen consumption rate (OCR) in young and aged control cells as well as after treatment with etomoxir. Etomoxir inhibits the CPT-1 enzyme and blocks FAO. Although etomoxir treatment of young cells only slightly decreased basal and maximal OCR, both basal and maximal OCR were significantly decreased upon etomoxir treatment in aged MSCs (Figure 3G and 3H), suggesting that aged cells up-regulate FAO to produce energy. Such a metabolic switch should be also reflected in the NAD+/NADH ratio. Indeed, we found that the NAD+/NADH ratio declined with age (Figure 3I). Together, our results demonstrate that during ageing, MSCs undergo a metabolic shift from glycolysis towards FAO, which together with the impaired fatty acid biosynthesis are responsible for the elevated acetyl-CoA levels. To further validate that the higher acetyl-CoA in the aged cells is indeed a result of its lower consumption during lipogenesis and its higher production during FAO, we focused on the different pathways through which acetyl-CoA is generated (Figure S3E). We compared ACLY and AceSC1 protein levels between young and aged cells, in order to investigate whether the different levels of acetyl-CoA are due to differential regulation of these enzymes. Both ACLY and AceCS1 levels were unaffected during ageing (Figure S3F), indicating that the conversion of citrate and acetate in the cytoplasm are not responsible for the observed changes in the acetyl-CoA levels. Therefore, the age-dependent metabolic alterations described above explain the observed increase in acetyl-CoA levels. However, they do not explain the apparent contradiction to the histone hypo-acetylation.

### Citrate carrier links mitochondrial acetyl-CoA pool with histone acetylation and stemness of MSCs upon ageing

As discussed above, histone acetylation relies on the local availability of acetyl-CoA (Wellen et al. 2009; Cai et al. 2011). Since aged cells contain higher levels of acetyl-CoA, one likely explanation for the reduced histone aceylation upon ageing could be the differential expression of histone acetyl-transferases (HATs). However, protein levels of CBP, which is a major HAT, remain unaffected upon ageing (Figure S4A). Given that the majority of acetyl-CoA is generated inside mitochondria as part of the TCA cycle, we speculated that the discrepancy between acetyl-CoA and histone acetylation levels might be due to decreased export of acetyl-CoA from the mitochondria to the cytoplasm. To test this hypothesis, we took into consideration the fact that inside mitochondria proteins are acetylated in a non-enzymatic manner, when acetyl-CoA levels are high (Hong et al. 2016; James et al. 2017).

We thus performed an immunofluorescence experiment and stained young and aged cells using an antibody against pan-acetyl-lysine, whereas TOMM20 was used as a counterstain for mitochondria. We observed a strong, age-dependent change in the localization of lysine acetylation signal, shifting from nuclear to mitochondrial upon ageing (Figure 4A).

**Figure 4.**
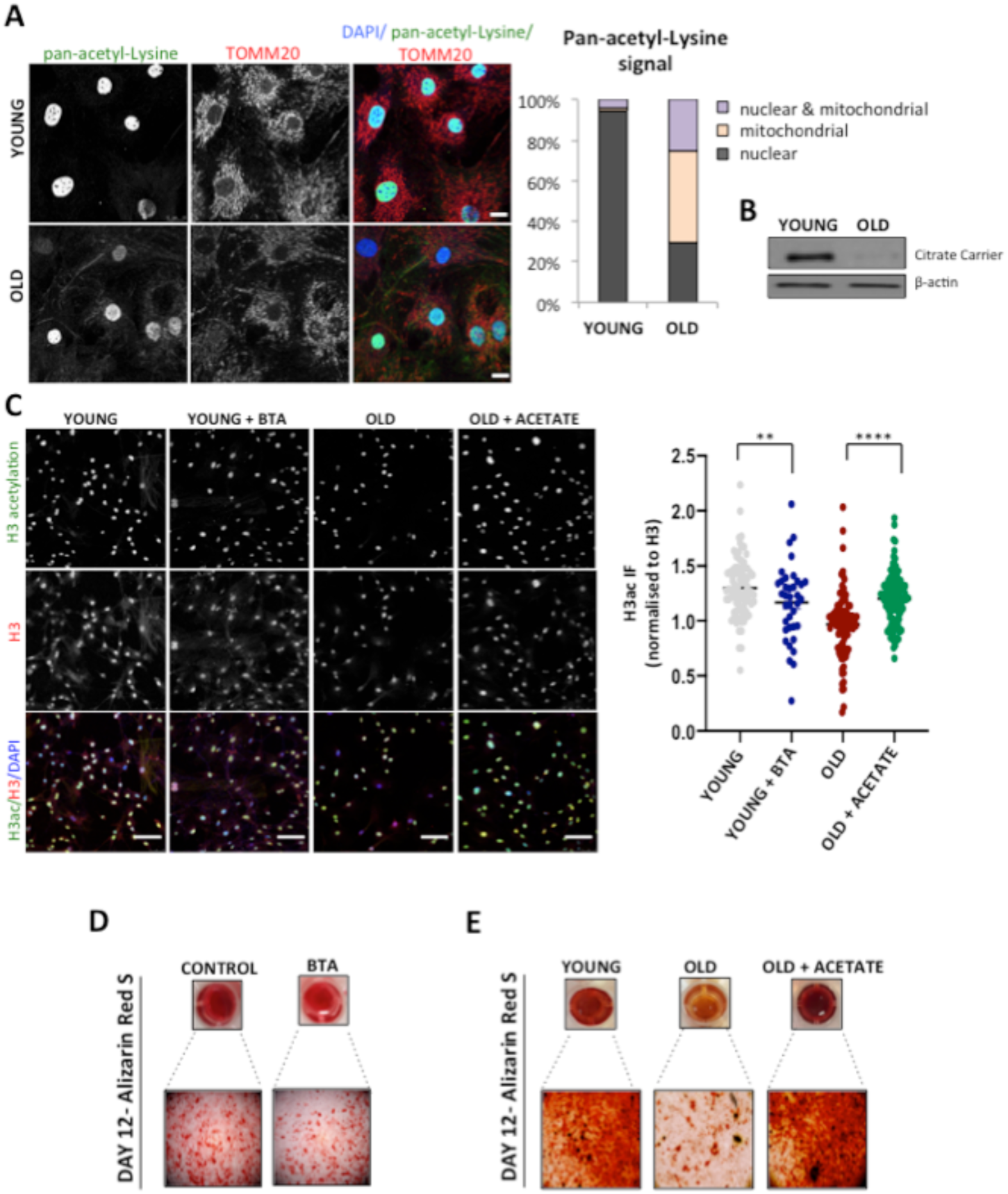
Citrate carrier links acetyl-CoA to histone acetylation and stemness upon ageing. **(A)** Representative images of young and aged MSCs after immunofluorescence staining, using pan-acetyl-Lysine and TOMM20 antibodies (left). Quantification of the localization of pan-acetyl-Lysine signal (right). Nuclei were stained with DAPI. Scale bars, 10μm. **(B)** Representative immunoblots for the citrate carrier protein in young and aged MSCs. β-was used as a loading control; (n=3 for each age group). **(C)** Representative images and quantification after immunofluorescence staining of young cells treated with or without 1mM BTA for 3 days, using antibodies against histone H3 acetylation and of aged cells treated with or without 5mM sodium acetate for 3 days. Levels of Histone H3 fluorescence intensity were used as an internal control, for normalization. Nuclei were stained with DAPI. Scale bars, 75μm. **(D)** Representative images of Alizarin-Red-S staining performed on young MSCs, 12 days after induction of osteogenesis. Cells were treated with or without 1mM BTA during the osteogenesis period; (n=3 biological replicates for each group). **(E)** Representative images of Alizarin-Red-S staining performed on young and aged MSCs, 12 days after induction of osteogenesis. Cells were treated with or without 5mM acetate for 3 days, before the induction of osteogenesis; (n=3 biological replicates for each group). Images were acquired using bright-field microscopy. Data are presented as mean ± SEM. Statistical significance was determined using student’s t-Test; **** p-value <0,001. See also Figure S4.

This suggests that acetyl-CoA is trapped inside the mitochondria of aged cells. To explain why aged cells showed such a compartmentalized localization of acetyl-CoA, we focused on the protein responsible for acetyl-CoA export. Citrate carrier (CiC) resides in the inner mitochondrial membrane and functions as an antiporter; it imports malate from the cytosol into mitochondria and exports citrate from mitochondria to the cytosol (Gnoni et al. 2009). In the cytosol, citrate is converted into acetyl-CoA by ACLY enzyme. Importantly, a connection between citrate carrier and histone acetylation has been demonstrated in *D. melanogaster* and in primary human fibroblasts (Morciano et al. 2009). Taking this into account, we speculated that down-regulation of CiC during ageing might result in lower cytoplasmic/nuclear levels of acetyl-CoA, explaining the observed decrease in histone acetylation. To address this possibility, we compared CiC mRNA and protein levels between young and aged MSCs. Strikingly, although mRNA levels of the *Slc25a1* gene which encodes the citrate carrier were only slightly affected upon ageing (Figure S4B), CiC protein levels were dramatically decreased in aged cells (Figure 4B), confirming our hypothesis that MSCs from old mice display impaired export of acetyl-CoA from mitochondria to the cytosol.

Next, we critically tested whether CiC was indeed the mechanistic target linking mitochondrial acetyl-CoA to histone acetylation in aged cells. Since the extremely limited amount of purified MSCs does not allow us to perform transfection-based studies, we further assessed the CiC-histone acetylation connection using two independent approaches. On one hand, we used 1,2,3-benzene-tricarboxylic acid (BTA), a specific inhibitor of the citrate transporter (Kolukula et al. 2014; Hlouschek et al. 2018), to treat young cells. After three days of treatment, we analyzed levels of histone H3 acetylation by immunofluorescence, as described above. Inhibition of CiC activity in young cells decreased levels of histone acetylation (Figure 4C), mimicking the results observed in aged MSCs. In a complementary approach, we added sodium acetate directly to the media of aged cells. Acetate represents an exogenous source of acetyl-CoA and can be converted into acetyl-CoA in the cytoplasm by AceCS1 enzyme (Moussaieff et al. 2015). This treatment would circumvent the impaired acetyl-CoA export from mitochondria to the cytosol. Notably, as shown in Figure S3F, AceCS1 levels were comparable between young and aged cells. Surprisingly, supplementation of aged cells with sodium acetate for three days rescued the loss of histone acetylation observed in the aged cells, to levels similar to those of young cells (Figure 4C). Collectively, these data demonstrate that loss of citrate carrier upon ageing is responsible for the reduction of histone acetylation levels, providing a mechanistic link between intermediate metabolism and histone acetylation.

We next sought to understand the biological consequences of the CiC-mediated mitochondrial-nuclear communication. The efficient differentiation of MSCs towards the adipogenic and osteogenic lineages requires oxidative metabolism and a fully functional lipid biosynthesis pathway (Wellen et al. 2009), including efficient export of the acetyl-CoA from mitochondria to the cytoplasm. Hence, we speculated that the decrease of citrate carrier levels observed in the aged cells might contribute to the lower differentiation potential of aged MSCs. To address this possibility, we treated young cells with BTA and we induced cell differentiation into adipocytes and osteocytes. Unexpectedly, although adipogenesis was only mildly affected by CiC inhibition (Figure S4C), osteogenic differentiation was strongly decreased in BTA-treated cells (Figure 4D). This behaviour of young MSCs treated with BTA mimics the behaviour of aged MSCs (Figure 1C and S1E), suggesting that CiC activity is required for successful osteogenic differentiation. To further confirm that the impaired acetyl-CoA export from mitochondria was responsible for the lower osteogenic differentiation capacity of MSCs from old mice, we supplemented media of aged MSCs with sodium acetate for three days and we then induced cell differentiation into osteocytes. Impressively, acetate treatment rescued the capacity of aged cells to give rise to osteocytes (Figure 4E), to levels similar to those in young cells. These data clearly demonstrate that citrate carrier mediates the connection between age-associated changes in chromatin, metabolism, and stemness upon ageing of MSCs.

## DISCUSSION

The data presented here suggest a model (Figure 5), whereby ageing-driven changes in metabolism and chromatin structure lead to transcriptional alterations and ultimately, to decreased osteogenic potential of aged MSCs. Our data demonstrate that the molecular link between mitochondrial metabolism and chromatin landscape is the citrate carrier. Upon ageing, lower citrate carrier levels result in impaired acetyl-CoA flux and thus de-regulated mitochondrial-nuclear communication and defective osteogenesis.

**Figure 5.**
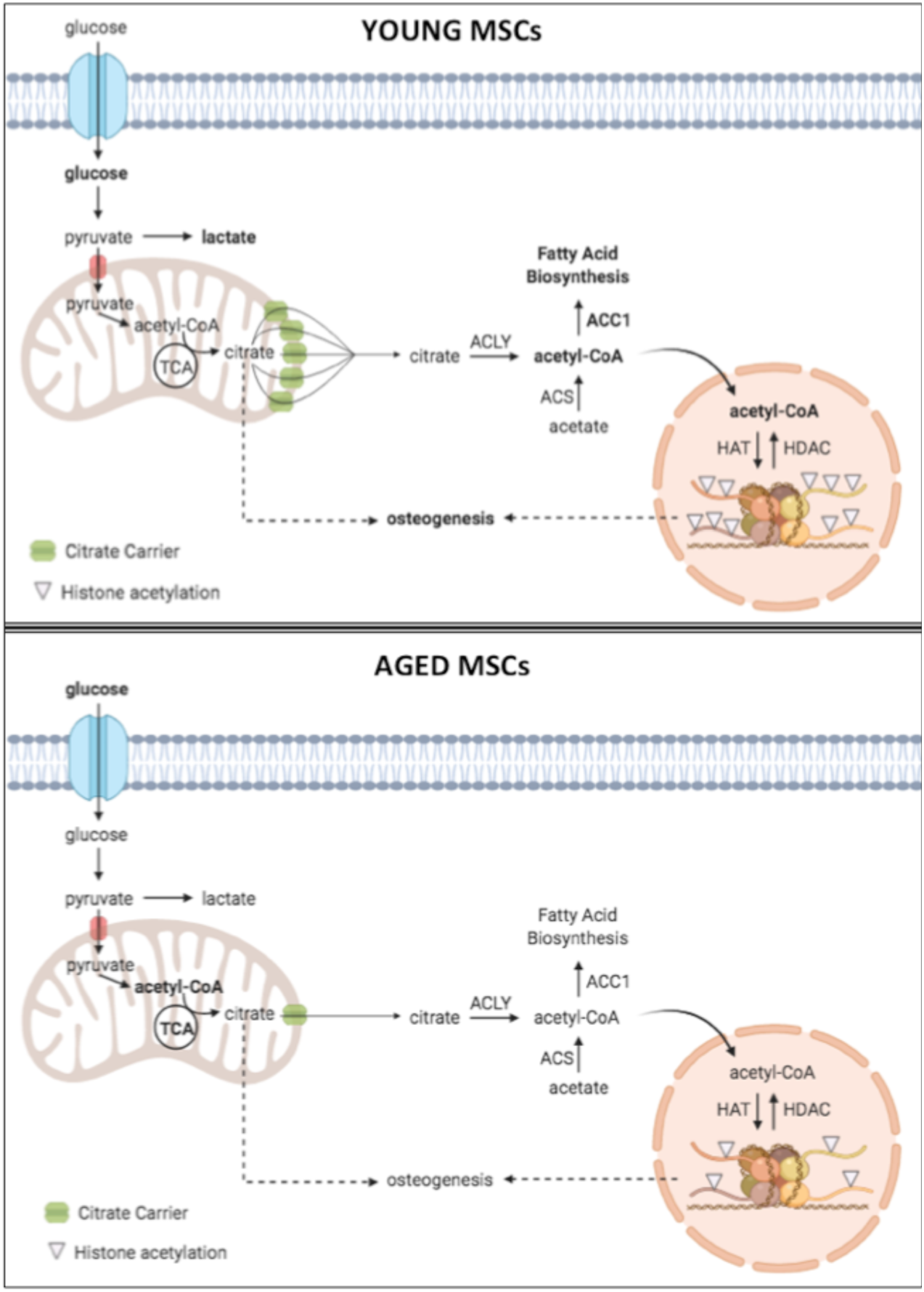
Model for citrate carrier-mediated connection among metabolism, chromatin and osteogenesis in MSCs. Upon ageing, MSCs perform glycolysis at a low extent, whereas they up-regulate fatty acid oxidation. This generates higher levels of mitochondrial acetyl-CoA. However, the cytosolic/nuclear acetyl-CoA pool of aged cells is decreased, due to lower citrate carrier levels. Concomitantly, aged cells exhibit decreased fatty acid biosynthesis and lower histone acetylation. The impaired export of acetyl-CoA from the mitochondria to the cytosol due to the reduced citrate carrier levels is responsible for the lower osteogenic potential of aged cells. Bold text indicates increase in accumulation and/or pathway.

We show that MSCs enriched for PDGFR? and Sca-1 markers exhibit chromatin compaction upon ageing. Interestingly, the age-associated effect on chromatin architecture appears to be cell- and tissue-type specific. For instance, human MSCs from peripheral blood show similar compaction of chromatin around gene promoters upon ageing (Ucar et al. 2017), whereas purified developing B cells hardly show any significant difference in chromatin accessibility (Koohy et al. 2018). It is crucial to highlight, though, that in contrast to physiological ageing, oncogene-induced senescence causes wide-spread changes in chromatin architecture with a strong, overall increase in chromatin accessibility (De Cecco et al. 2013; Parry et al. 2018). In addition to monitor global chromatin accessibility, we used our ATAC-seq dataset to predict changes in transcription factor activity during ageing. The most prominently enriched motif belongs to CTCF (Figure 2E), a key component of chromatin that is critical for 3D genome organization, transcription control and enhancer/promoter insulation (Braccioli and de Wit 2019). Although we do not have any further insight into the role of CTCF in the ageing process of MSCs, it is tempting to speculate that CTCF, as a major structural chromatin-associated protein, is part of the network that drives the observed changes in accessibility during ageing. Metabolic switches occur frequently during development as well as upon differentiation and a pluripotent stem cell state is usually strongly correlated with a specific metabolic profile (Shyh-Chang, Daley, and Cantley 2013). In particular, multipotent MSCs rely on glycolysis whereas they up-regulate oxidative phosphorylation upon differentiation into the adipogenic and osteogenic lineages. Here, we demonstrate that aged MSCs show lower glycolytic rate and rely more on FAO, which results in higher acetyl-CoA production (Figure 3). However, acetyl-CoA remains inside the mitochondria of aged cells and it cannot be used for acetylation of histones, due to lower citrate carrier levels (Figure 4). Importantly, non-enzymatic protein acetylation in mitochondria has been previously described (Hong et al. 2016; James et al. 2017) and acetylation of mitochondrial proteins might have broader consequences on their function, that might be worth investigating in the future.

As shown in Figures 1D and 1E, we found that the age-associated decrease in MSC differentiation capacity affects only the trabecular bone compartment, whereas the quality of the cortical bone remained unaffected upon ageing, consistent to previous reports (Ritzel et al. 1997; Eswaran et al. 2006; Christiansen et al. 2011). A likely explanation of this observation is that MSCs reside almost exclusively in the trabecular bone, thereby suggesting that this compartment is more heavily affected by defects in MSC function.

We identified here that citrate carrier links chromatin architecture and stem cell fate decisions upon ageing. However, it is important to stress that this interconnection might be cell type-specific. A recent study (Hennrich et al. 2018) used proteomic analysis to profile changes in the proteome of human bone marrow-resident cells, including hematopoietic stem cells and progenitor cells (HPCs), from 59 individuals between 20 and 60 years old. Interestingly, citrate carrier as well as several proteins of the initial glycolytic steps are up-regulated in aged HPCs, which is likely a reminiscence of the Warburg effect. The upregulation of the citrate carrier might also be linked to the higher proliferative capacity of HPCs compared to MSCs, with age (Akunuru and Geiger 2016). This phenomenon is also observed in cancer cells (Fernandez et al. 2018). In contrast to our observations, the authors did not report any change in citrate carrier levels in MSCs between donors of different ages. One possible explanation is the differences in the isolation protocol and the cell culture conditions. Hennrich et al. isolated MSCs following the classical plastic adherence approach (Wagner et al. 2008), which isolates a much more diverse set of MSCs. In addition, expansion of MSCs was done at atmospheric oxygen, whereas we cultured MSCs permanently under low oxygen concentration, more accurately mimicking their physiological niche. Given that oxygen tension has a great impact on various energy-producing pathways and in particular on mitochondrial metabolism, these differences in cell culture conditions might explain the differences in CiC levels.

Understanding the molecular mechanism mediating the mito-nuclear communication during ageing in MSCs enabled us to rescue the age-related osteogenic differentiation defect, by acetate supplementation of aged MSCs (Figure 4E). Collectively, our work identifies a single mitochondrial protein as a focal point for the regulation of metabolic homeostasis and stem cell potential upon ageing. These results highlight the tight connection between metabolic and chromatin states on the regulation of stem cell fate and propose citrate carrier as an attractive target for therapeutic intervention and *in vitro* rejuvenation of aged MSCs.

## ACKNOWLEDGEMENTS

We would like to thank all members of the Tessarz lab for discussion together with Martin Graef and Thomas Langer for critical reading of the manuscript. We are very grateful to the MPI AGE Comparative Biology facility for mouse housing. Sequencing was performed in the core facilities of the Max Planck Society at the Max Planck Institute for Plant Breeding Research (Cologne) and Molecular Genetics (Berlin). This work was funded by the Max Planck Society (to P.T. and L.P.), the NIH (to I.T and R.D., NIH GM125871), an Alexander von Humboldt postdoctoral fellowship to M.M. and an Onassis foundation graduate fellowship to A.P. Models were drawn using BioRender.

## AUTHOR CONTRIBUTION

Conceptualisation: A.P., M.M. and P.T.; Methodology: A.P., S.P., M.M. I.T., Y.H, K. F.-D., P.G. ; Investigation: A.P., M.M., M.B., C.N., K.F.-D., Y.H., A.M. and P.G.; Formal Analysis: S.P., I.T. and P.G.; Supervision: R.D., L.P. and P.T.; Project Administration and Funding Acquisition: P.T.; Writing of Manuscript: A.P. and P.T.

## DATA AVAILABILITY

ATAC- and RNA-seq data are available at GEO, accession code: GSEXXXX

## MATERIALS AND METHODS

### Antibodies

For cell sorting, CD140a-APC (#17140181), Sca-1-FITC (#11598185), Terr-119-PE (#12592182) and the Fixable Viability Dye eFluor 450 (#65086314) were all purchased from eBioscience, whereas CD45-PE (#A16325) antibody was obtained from Life Technologies. For immunofluorescence and Western Blot experiments, antibodies against pan-acetylated-Lysine (#9441S), Fatty Acid Synthase (#3189S), CBP (7389S), AceCS1 (#3658T), anti-rabbit IgG HRP-linked (#7074S), anti-mouse IgG HRP-linked (#7076S) and Histone H3 (#14269) were all purchased from Cell Signaling Technology. Antibodies against ACC1 (#21923-1-AP), ACLY (#15421-1-AP) were from ProteinTech. Antibodies against TOMM20 (sc-17764) and b-actin (sc-47778) were bought from Santa Cruz Biotechnology; another TOMM20 antibody (WH0009804M1) was bought from Sigma-Aldrich. Histone H3 pan-acetyl (#39139) antibody was obtained from Active Motif, whereas Histone H4 pan-acetyl (06-866) was bought from EMD Millipore. Anti-Mouse IgG Alexa fluor 488 (#A11001) and anti-Rabbit IgG Alexa Fluor 488 were bought from ThermoFisher Scientific. Citrate carrier antibody (#99168) and Histone H4 antibody (#31830) were obtained from Abcam.

## MOUSE MODEL AND EXPERIMENTAL DETAILS

### Mice

C57BL/6N mice were bred and cared for in the mouse facility of Max Planck Institute for Biology of Ageing following ethical approval by the local authorities. For our ageing experiments and profiling we used exclusively wild type male mice at 3-4 months (young cohort) and 18-22 months (old cohort) of age, a range commonly used in age-associated studies.

### Endosteal-MSCs isolation

To isolate MSCs from their endosteal niche we followed a purification strategy based on a published protocol (Houlihan et al., 2012), and adapted the isolation strategy to acquire sufficient cells for our experiments. In brief, young and old C57BL6/N mice were sacrificed by cervical dislocation. Skin and muscles around the hind limbs were removed, the legs were cut above the pelvic joints and then placed in ice-cold PBS, on ice. Tibias were then separated from femurs by dislocating the joints and the clean bones were placed back in ice-cold PBS. All the following steps were performed inside the hypoxia hood. Bones were crushed and cut into tiny pieces and bone chips were incubated at 37°C for 75 minutes, in a-MEM medium containing 0.2% w/v collagenase (Sigma Aldrich), shaking at 200 rpm. To stop collagenase reaction, sample tubes with the bone chips were placed on ice and washed with a-MEM medium supplemented with 10% FBS and 1% penicillin/streptomycin. We modified the published protocol by culturing the bone fragments together with the released cells; this allows the outgrowth of more MSCs from the bone *in vitro* before cell sorting by flow cytometry, increasing the cell yield we obtain. Bone chips were transferred into T25 flasks and were cultured under humidified conditions in hypoxia (2% O_2_, 5%CO_2_, 37°C). On day 3 of cell culture, the medium was changed and on day 5 both the cells and the bone chips were passaged, using Trypsin/EDTA solution (Life Technologies). On day 8 of the cell culture the bone chips were removed and on day 12 we performed cell sorting using flow cytometry.

### Cell sorting by Flow Cytometry

To obtain a purified MSC population we performed cell sorting by flow cytometry, using Fixable Viability Dye eFluor 450 (1:10000) and the following antibodies: Sca-1-FITC, CD140a-APC, CD45-PE and TER-119-PE, all in 1:1000 dilution. After harvesting, the cells were washed with PBS and resuspended in Hank’s Balanced Salt Solution (1x HBSS, 0.01M HEPES, 2% FBS, 1% penicillin/streptomycin, all Life Technologies). After incubation with the antibodies for 45 minutes, cells were washed twice with HBSS+ and were filtered through 35μm nylon mesh into 5ml sample tubes. Sorting was performed using BD FACSAria IIu and BD FACSAria IIIu instruments, under low pressure sorting conditions (100μm nozzle and 20 psi sheath pressure). The CD45-/TERR-119-/CD140a+/Sca-1+ population was sorted into an eppendorf tube containing 500μl a-MEM medium, at 4°C. Compensation was done using UltraComp compensation beads (#01222242, Invitrogen,) and ArC beads (#A10346, Life technologies). Once sorted, MSCs were centrifuged at 300g for 10 minutes at 4°C. They were then resuspended in a-MEM medium, supplemented with 10% FBS and 1% penicillin/streptomycin. Cells were cultured under humidified conditions in hypoxia (2% O_2_, 5%CO_2_, 37°C), in T25 flasks.

### qRT-PCR

Cells were lysed with QIAzol (QIAGEN) and total RNA was extracted using RNA extraction kit (Direct-zol RNA MiniPrep - Zymoresearch), following the manufacturer’s protocol. This was followed by cDNA synthesis using Maxima™ H Minus cDNA synthesis master mix (Thermo Scientific), according to the manufacturer’s instructions. Subsequent qRT-PCR was performed with 10ng of cDNA, using SYBR-Green chemistry (Roche) on a Light Cycler 96 instrument (Roche). Data were analyzed and further processed in Microsoft Excel and Prism8 software. Fold change in gene expression over control samples was calculated using the ΔΔCq method, where β-actin Cq values were used as internal control. All reactions were run in three technical replicates and averaged.

Oligos were designed using Primer3 and Blast platforms. The primers used for each gene are:

#### Slc25a1 (encoding citrate carrier)

5’ GGAGAGGACTATTGTGCGGTCT 3’

5’ CCCGTGGAAAAATCCTCGGTAC 3’

#### Acan

5’ CGTTGCAGACCAGGAGCAAT 3’

5’ CGGTCATGAAAGTGGCGGTA 3’

#### β-actin

5’ CTGCGCTGGTCGTCG 3’

5’ CACGATGGAGGGGAATACAG 3’

### Measurement of mtDNA content

RT-qPCR of mitochondrial DNA-encoded genes was used to estimate the mitochondrial DNA content. Cells were lysed with DNA lysis buffer (0.5% SDS, 0.1M NaCl, 50mM Tris pH=8.0, 2.5mM EDTA pH=8.0) and lysates were incubated on a Thermoshaker (620 rpm, 55°C) for 30 minutes, with Proteinase K (Sigma Aldrich). Phenol (Roth) was then added to the samples to denature proteins and after this, equal amount of chloroform (Roth) was added to remove residual phenol. To precipitate DNA, the aqueous phase was collected again and samples were incubated overnight at -80°C with sodium acetate (pH= 5.2; Invitrogen and 100% ethanol (Roth). Precipitated DNA was then washed with 70% ethanol for 20 minutes and after airdrying, it was resuspended in nuclease free water.

RT-qPCR for atp6 and cox1 genes was performed using SYBR-Green chemistry with the Roche Light Cycler 96 instrument, as described above. Data were analyzed and further processed in Microsoft Excel and Prism8 software. Fold change in gene expression over control samples was calculated using the dCt method and genomic β-actin Ct values were used for internal normalization. All reactions were run in three technical replicates and averaged.

Oligos were designed as above. The primers used for each gene are:

#### atp6

5’ GGCACCTTCACCAAAATCAC 3’

5’ CGGTTGTTGATTAGGCGTTT 3’

#### cox-1

5’ AGGTTGGTTCCTCGAATGTG3’

5’ GCCTTTCAGGAATACCACGA 3’

### Mitotracker staining

Cells were washed with PBS, harvested with Trypsin-EDTA (Life Technologies) and resuspended in the pre-warmed (37°C) staining solution containing the MitoTracker Deep Red FM probe (ThermoFischer Scientific #M22426) in a final dilution 1:15000 (in a-MEM medium without FBS and phenol red). Cells were incubated with the staining solution for 30 minutes at 37°C, under hypoxic conditions. They were then washed twice with PBS, and centrifuged at 500g for 5 minutes. In the end, they were resuspended in a-MEM medium without FBS and phenol red. After addition of DAPI (Invitrogen) for dead-cell exclusion right before measurement, cells were analyzed by flow cytometry (BD FACSCANTO II cytometer, BD Biosciences). Data were collected using FACS-Diva software and analyzed using FlowJo software.

### Measurement of bone parameters by μCT

To assess changes in bone quality upon ageing, we dissected femurs from young and old mice and collected them in PBS. Mouse femurs were then scanned with a high resolution µCT scanner (SkyScan 1176, Bruker, Belgium) with an isotropic voxel size of 8.8 µm3. The x-ray settings for each scan were 50 kV and 200 µA using a 0.5 mm aluminum filter. All scans were performed over 360 degrees with a rotation step of 0.3 degrees and a frame averaging of 1. Images were reconstructed and analyzed using NRecon and CTAn software, respectively (Bruker, Belgium). Trabecular and cortical bone regions of distal femurs were selected with reference to the growth plate (0.44-2.2 and 2.2-2.64 mm from growth plate, respectively). Bone mineral density was determined based on calibration with two phantoms of known density (Bruker, Belgium), which were scanned under the same conditions as the bone samples.

### Oxygen Consumption

The SeaHorse XF96 extracellular Flux Analyzer (Agilent Technologies) was used to determine oxygen consumption rate in young and aged cells. 20000 cells were seeded in 96-well SeaHorse plates, after coating them with 10% Gelatin - 90% Poly-L-Lysine solution for 1 hour. Cells were incubated overnight at 37°C, in a humidified 5% CO_2_, 2% O_2_ incubator, with 200μl of a-MEM medium, supplemented with 10% FBS and 1% Penicillin/Streptomycin. On the day of the experiment cells were treated with or without 100μM etomoxir (Sigma Aldrich) and incubated for 1 hour at 37°C, in a humidified 5% CO_2_, 2% O_2_ incubator. They were then washed twice with assay medium (XF-DMEM medium supplemented with 10mM Glucose, 1mM Pyruvate and 2mM L-Glutamine) and incubated with this medium for 1 hour prior to loading into the XF Analyzer, in a non-CO_2_ -containing incubator. Following measurements of resting respiration, cells were treated subsequently with 20μM oligomycin, 5μM FCCP and 5μM Rotenone/Antimycin (all from Agilent Technologies). Each measurement was taken over a 2-min interval followed by 2-min of mixing and 2-min of incubation. Three measurements were taken for the resting OCR: after oligomycin treatment, after FCCP and after Rotenone/Antimycin A treatment. Values were normalized to protein concentration using Bradford kit and were plotted using the Wave 2.4 and the Prism8 software.

### Colony-Forming Unit (CFU) assay

To determine the efficiency by which MSCs form colonies 10^2^ cells were seeded in a 6-well plate and cultured for 10-12 days at 37°C, in a humidified 5% CO_2_, 2% O_2_ incubator. Plates were washed with PBS and stained with 1% (v/v) Crystal Violet solution (Sigma Aldrich) for 5-10 minutes at room temperature. Cells were then washed thoroughly with water for 3 times and visible colonies were counted. Images were acquired on a CanonScan 9000F Mark II scanner.

### Differentiation assays

#### i) Differentiation to adipocytes

For adipogenesis experiments, 2×10^3^ cells were seeded in 96-well plates. Adipogenic differentiation was induced once cells reached confluency, by culturing them in control (a-MEM medium supplemented with 10% FBS and 1% penicillin/streptomycin) or differentiation (a-MEM medium supplemented with 10% FBS, 1% penicillin/streptomycin and i) 1μM dexamethasone (Sigma Aldrich) ii) 1μM IBMX (Sigma Aldrich) iii) 10μg/ml insulin (Sigma Aldrich) and iv) 100μM indomethacin (Sigma Aldrich)) media, for 8-10 days. For BTA treatment of young cells, 1mM BTA (Sigma Aldrich) was added in the control and adipogenic media throughout the differentiation period.

Adipocytes were detected by Oil-Red-O (Sigma Aldrich) staining. Cells were washed with PBS and fixed in 3.7% formaldehyde (Roth) for 30 minutes, at room temperature. After fixation, cells were washed twice with ddH_2_O and once with 60% isopropanol (Roth), with every washing step lasting 5 minutes. Cells were then stained with Oil-Red-O staining solution for 15 minutes at room temperature. After staining, cells were washed four times with ddH_2_O and images were acquired with bright-field microscope, using the 20x objective.

#### i) Differentiation to osteoblasts

For osteogenesis experiments, 2×10^3^ cells were seeded in 96-well plates. Osteogenesis was induced once cells reached confluency, by culturing them in control (a-MEM medium supplemented with 10% FBS and 1% penicillin/streptomycin) or differentiation (a-MEM medium supplemented with 10% FBS and 1% penicillin/streptomycin and i) 100nM dexamethasone ii) 10mM beta-glycerophosphate (Sigma Aldrich) iii) 100 μM ascorbic acid (Sigma Aldrich)) media, for 11 days. For BTA treatment of young cells, 1mM BTA was added in the control and osteogenic media throughout the differentiation period. For acetate treatment of aged cells, 5mM of sodium acetate was added in the media three days prior to the induction of differentiation and was then removed.

Osteoblasts were detected by Alizarin Red S (Sigma Aldrich) staining, performed 12 days after induction of osteogenesis. Cells were washed once with PBS and fixed in 3.7% formaldehyde for at least 30 minutes, at room temperature. Fixation was followed by washing of cells with ddH_2_O. Cells were then incubated with the Alizarin Red S staining solution (2% w/v Alizarin Red S in ddH_2_O) for 45 minutes, protected from light. After staining, cells were washed with ddH_2_O and images were acquired with bright-field microscope, using the 20x objective.

#### i) Differentiation to chondrocytes

For chondrogenic experiments, 75×10^3^ cells were seeded in 96-well plates with flat (control) or conical (differentiated) bottom. Chrondrogenesis was induced once cells reached confluency, by culturing them in control (a-MEM medium supplemented with 10% FBS and 1% penicillin/streptomycin) or differentiation (DMEM medium supplemented with 10% FBS and 1% penicillin/streptomycin and i) 100nM dexamethasone ii) 1% ITS (SIgma Aldrich), iii) 10 μM ascorbic acid iv) 1mM sodium pyruvate (Gibco) v) 50 μg/ml proline (SIgma Aldrich) and vi) 20nm/ml TGFb3 (SIgma Aldrich)) media for 12-14 days.

Confirmation of osteogenesis was performed by extracting RNA and running qPCR analysis for osteogenesis markers, as described above.

### Immunofluorescence experiments

For immunofluorescence experiments, 2×10^3^ cells were seeded in 96-well plates with glass bottom (Greiner) and treated as indicated in each experiment. After treatments, cells were fixed for 15 minutes at 37°C with 3.7% v/v formaldehyde in a-MEM medium, supplemented with 10% FBS and 1% penicillin/streptomycin. Samples were washed with PBS twice, permeabilized with 0.1% TritonX-100 (Roth) in PBS for 10-15 minutes and blocked with 5% BSA in PBS (Roth) for 10 minutes. Samples were then incubated with the indicated primary antibodies in 5%BSA-PBS, in dilutions 1:100-1:300 overnight at 4°C. Following this, they were washed three times with PBS, with each washing step lasting 10 minutes. Samples were then incubated with the appropriate secondary fluorescent antibodies diluted 1:500 for 45 minutes, protected from light. After three washing steps of 10 minutes each, cells were mounted using Roti-Mount FluorCare mounting medium (HP20.1, ROTH), containing DAPI. Images were acquired using 40x and 100x objective lens on SP8X leica confocal microscope.

### Western Blot experiments

For all western blot experiments cells were harvested and lysed with RIPA buffer (150mM NaCl, 1% TritonX-100, 0.5% sodium deoxycholate, 0.1% SDS, 50mM Tris pH=8.0) supplemented with 5mM sodium butyrate and 1x Protease Inhibitor Cocktail (Thermo Scientific). For efficient lysis, cells were incubated with RIPA buffer at 4°C, for 30 minutes, rotating, and then centrifuged for 10 minutes at 6500g. Protein concentration was determined using BCA protein assay kit (SERVA). 20-50μg of total protein were loaded into each well and SDS-PAGE electrophoresis was performed at 160V for ∼45 minutes. This was followed by transfer to nitrocellulose membrane, using the Trans-Blot Turbo blotting apparatus and reagents, all provided by Bio-Rad. Protein transfer was confirmed by Ponceau S (Sigma Aldrich) staining for 1-2 minutes. The membranes were then blocked using 5% non-fat dry milk in Tris-buffered saline-0.1% Tween20 (TBS-T) for 1 hour at room temperature (RT). Membranes were incubated with the indicated primary antibodies, diluted in 3% Milk in TBS-T, at 4°C overnight, washed three times with PBS for 5 minutes each washing step and incubated with the appropriate horseradish peroxidase (HRP)-conjugated secondary antibodies diluted 1:10000 in 5% BSA in TBS-T, for 1 hour at RT. After three 10-minute washing steps in TBS-T, the desired proteins were visualized by providing fresh HRP-substrate solution (Luminol Enhancer Solution/Peroxide Solution - Promega) and exposure of membranes for specific time periods to photographic film, using the Curix60 instrument (Agfa).

### Measurement of glucose, lactate and pH

Measurement of glucose, lactate and pH in the media of MSCs was performed using the Vi-CELL MetaFLEX instrument (Beckman Coulter), according to manufacturer’s instructions. All measurements were done in triplicates and averaged.

### Metabolite extraction for Liquid Chromatography mass spectrometry (LC-MS)

Cells were cultured in a-MEM medium supplemented with 10% FBS and 1% penicillin/streptomycin, harvested using Trypsin-EDTA (Life Technologies) and snap-frozen in liquid nitrogen. Metabolite extraction from each cell pellet was performed using 1 mL of a mixture of 40:40:20 [v:v:v] of pre-chilled (−20°C) acetonitrile:methanol:water (OptimaTM LC/MS grade, Thermo Fisher Scientific). The samples were subsequently vortexed until the cell pellets were fully suspended, before incubating them on an orbital mixer at 4°C for 30 minutes at 1500 rpm. For further disintegration, samples were sonicated for 10 minutes in an ice cooled bath sonicator (VWR, Germany) before centrifuging them for 10 minutes at 21100x g and 4°C. The metabolite-containing supernatant was collected in fresh tubes and concentrated to dryness in a Speed Vac concentrator (Eppendorf). The protein-containing pellets were also collected and used for protein quantification (BCA Protein Assay Kit, Thermo Fisher Scientific).

#### i) Targeted LC-MS analysis of acetyl CoA, NAD and <u>NADH</u>

For the analysis of acetyl-CoA, NAD+ and NADH the extracted metabolites were re-suspended in 40 µl of UPLC-grade acetonitrile:water (80:20 [v:v], OptimaTM LC-MS-grade, Thermo Fisher Scientific). The samples were analyzed on an Acquity iClass UPLC (Waters), using a SeQuant ZIC-HILIC 5µm polymer 100 × 2.1 mm column (Merck) connected to a Xevo TQ-S (Waters) triple quadrupole mass spectrometer.

8µl of the re-suspended metabolite extract were injected onto the column and separated using a flow rate of 500 µl/minute of buffer A (10mM ammonium acetate, 0.1% acetic acid) and buffer B (acetonitrile) using the following gradient: 0 - 0.5 minutes 20% A; 0.5 - 1.4 minutes 20 - 35% A; 1.4 – 2.5 minutes 35 - 65% A. After 2.5 minutes the system is set back to 20% A and re-equilibrated for 2.5 minutes.

The eluting metabolites were detected in positive ion mode using ESI MRM (multi reaction monitoring) applying the following settings: capillary voltage 1.5 kV, desolvation temperature 550°C, desolvation gas flow rate 800 L/h, collision cell gas flow 0.15 mL/min. The following MRM transitions were used for relative compound quantification of acetyl CoA m/z precursor mass (M+H+) 810, fragment mass (M+H+) m/z 303 using a cone voltage of 98V and a collision energy of 28V, NAD m/z precursor mass (M+H+) 664, fragment mass (M+H+) m/z 136 using a cone voltage of 50V and a collision energy of 56V and NADH m/z precursor mass (M+H+) 665, fragment mass (M+H+) m/z 108 using a cone voltage of 70V and a collision energy of 70V. For each compound two additional fragments were monitored as qualitative controls for compound identity. Data analysis and peak integration were performed using the TargetLynx Software (Waters).

### Gas Chromatography mass spectrometry (GC-MS) for the analysis of TCA cycle intermediates and amino acids

High resolution GC-MS analysis (Q-Exactive GC, Thermo Fisher Scientific) was performed for the analysis of amino acids and metabolites from the TCA cycle. For this purpose, the dried metabolite extracts were derivatized using methoxyamine (methoxyamine hydrochloride, Sigma) and N-Methyl-N-trimethylsilyl-trifluoracetamid (MSTFA, Macherey-Nagel) before performing the GC-MS analysis. In brief, dried samples were resuspended in a freshly prepared (20 mg/mL) solution of methoxyamine in pyridine (Sigma). The re-suspended pellets were incubated for 90 minutes at 40°C on an orbital shaker (VWR) at 1500 rpm. After adding additional 45 µL of MSTFA, the samples were incubated for an additional 30 minutes at 40°C and 1500 rpm. At the end of the derivatization the samples were centrifuged for 10 minutes at 21100x g and 40 µL of the clear supernatant was transferred to fresh autosampler vials with conical glass inserts. For the GC-MS analysis 1 µL of each sample was injected using a PAL autosample system (Thermo Fisher Scientific) using a Split/Splitless (SSL) injector at 300 C in splitless mode. The carrier gas flow (helium) was set to 2 ml/min using a 30m DB-35MS capillary column (0.250 mm diameter and 0.25 µm film thickness, Agilent). The GC temperature program was the following: 2 minutes at 85°C, followed by a 15°C per minute ramp to 330°C. At the end of the gradient, the temperature was held for additional 6 minutes at 330°C. The transfer line and source temperature were both set to 280°C. The filament, which was operating at 70V, was switched on 2 minutes after the sample was injected. During the whole gradient period the MS was operated in full scan mode covering a mass range m/z 70 and 800 with a scan speed of 20 Hertz.

The GC-HRMS data were analysed and quantified using the TraceFinder (Thermo Fisher Scientific) software tool. The identity of each compound was validated by authentic reference compounds, which were analysed in independent GC-MS runs.

### RNA-seq

Total RNA was isolated using the RNA extraction kit (Direct-zol RNA MiniPrep - Zymoresearch), following the manufacturer’s protocol. Once RNA quality and integrity were verified, RNA libraries were created and the desired cDNA fragments were sent for sequencing. Libraries were sequenced as single-end 150bp reads on Illumina HiSeq 4000. The sequenced reads of RNA-seq dataset were processed using zUMIs (version 2.2.1) (Parekh et al. 2018) with STAR (version 2.6.1a) (Dobin et al. 2013), samtools (version 1.9) (H. Li et al. 2009) and featureCounts from Rsubread (version 1.32.4) (Liao, Smyth, and Shi 2014). The reads were mapped to the mouse genome (mm10) with the ensembl annotation verion GRCm38.91. The generated count matrix was further analysed using R (version 3.5.1). First, genes were filtered using “filterByExpr” function of edgeR (Robinson, McCarthy, and Smyth 2010) with the min.count=5. The differential gene expression analysis between young and old mice was carried out using limma-trend (Ritchie et al. 2015) approach at the adjusted p-value of 0.05. Obtained sets of genes were further analysed, e.g. through gene ontology (GO) enrichment analysis.

### ATAC-seq

The fastq files of sequenced reads were mapped to the mouse genome (mm10) using local alignment within bowtie2 (Langmead and Salzberg 2012) with parameters -x mm10 and -X 2000. The resulting BAM files were sorted, indexed using samtools (version 1.3.1) and duplicates were removed using MarkDuplicates of Picard Tools.

### DAStk analysis

The open chromatin regions were called from the resulting BAM files from mono-nucleosome, di-nucleosome, tri-nucleosome, and nucleosome-free regions using MACS2, with settings -- shift -100 --extsize 200 -B --nomodel --format BAMPE. Any ATAC-seq peaks overlapping blacklisted regions for the mm10 mouse reference genome (ENCODE ENCFF547MET) were excluded from the analysis.

We used mouse motifs from the HOCOMOCO (Kulakovskiy et al. 2018) database version 11, and scanned for coordinates of motif sites in the mm10 mouse reference genome utilizing FIMO(Grant, Bailey, and Noble 2011) at a p-value threshold of 10-6. DAStk (https://github.com/Dowell-Lab/DAStk) was utilized to predict changes in transcription factor activity, using a p-value < 0.05 cutoff for statistical significance.

## SUPPLEMENTARY FIGURES

**Figure S1.**
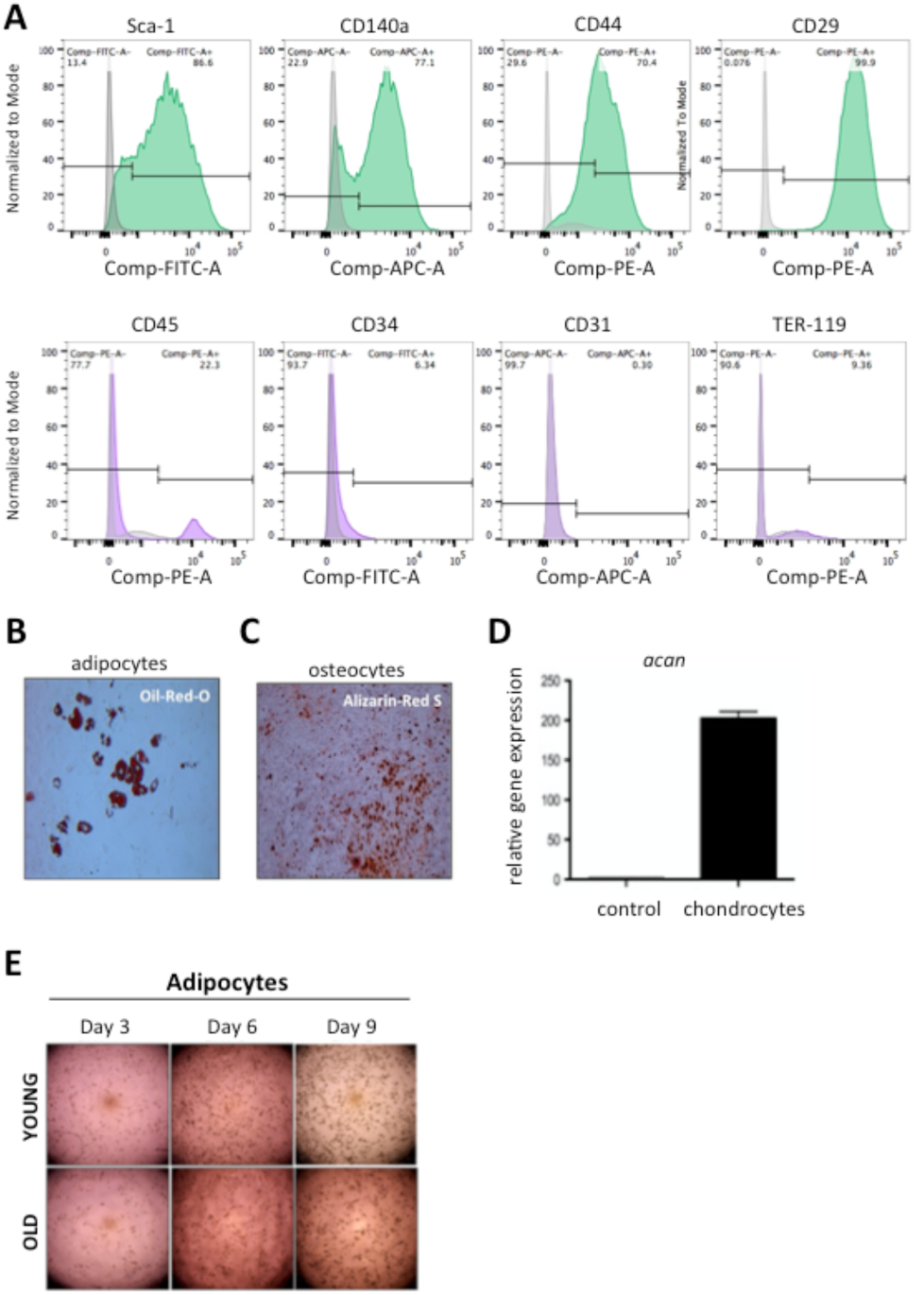
(related to Figure 1) **(A)** Expression profile of MSCs for mesenchymal (top) and hematopoietic (bottom) markers, as assessed by FACS analysis. Our purified MSCs population is enriched for mesenchymal stem cell markers and depleted for hematopoietic stem cell markers. **(B-D)** Multi-lineage differentiation capacity of MSCs. Representative images showing adipogenic and osteogenic potential, which were confirmed with Oil-Red-O and Alizarin-Red-S staining of MSCs, respectively. Images were acquired by bright-field microscopy, 12 days after induction of differentiation. Chondrogenic differentiation was confirmed by qPCR analysis of the chondrogenic gene *Acan*, 14 days after induction of differentiation and in comparison to non-induced cells. β-actin was used as an internal control for normalization (n=3 biological replicates in each experiment). **(E)** Bright-field microscopy images of adipocytes in young and aged cells 3 days, 6 days and 9 days after induction of adipogenesis.

**Figure S2.**
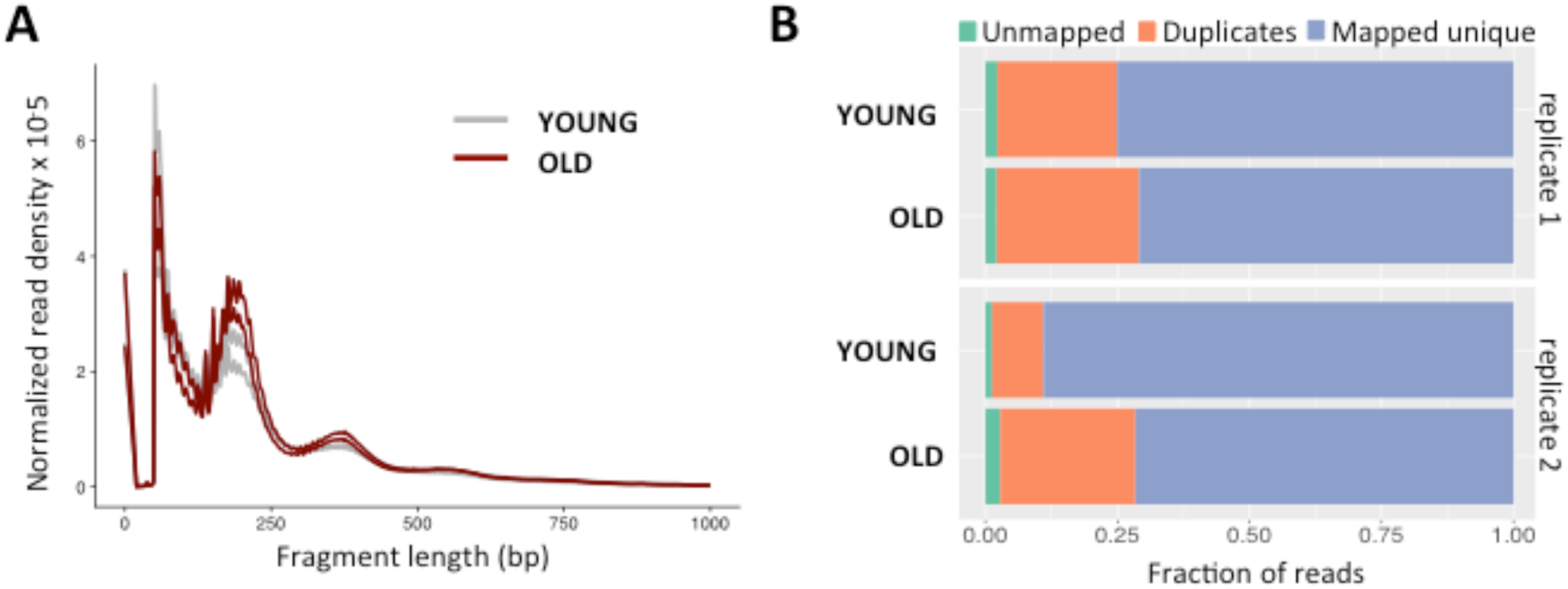
(related to Figure 2) Quality measures for ATAC-seq. **(A)** Insert size distribution of each of the four libraries (2 per age group) and **(B)** Mapping statistics.

**Figure S3.**
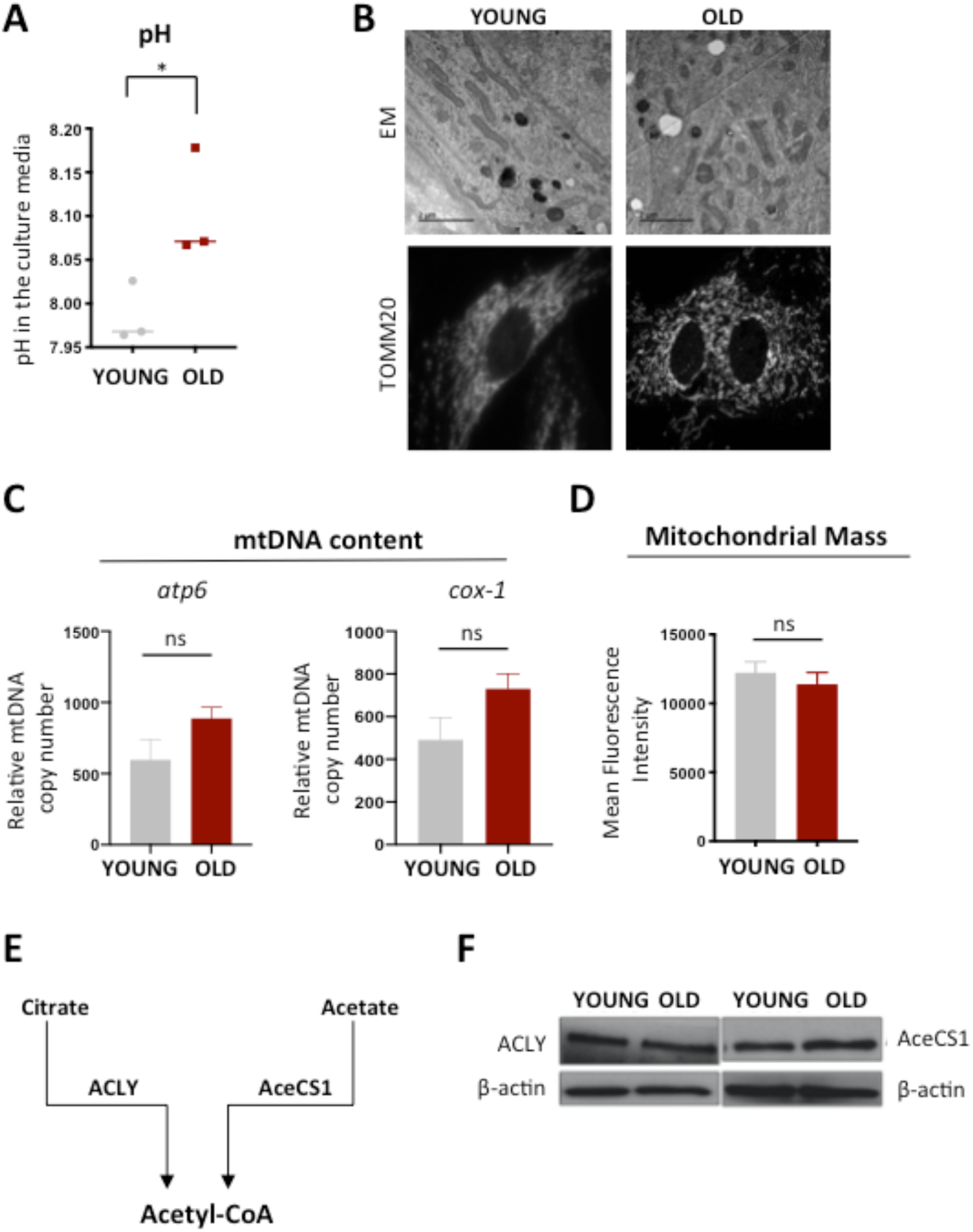
(related to Figure 3) **(A)** pH measurement of a-MEM media from young and aged cells, using the Vi-Cell MetaFLEX instrument; (n=3 biological replicates for each age group). **(B)** Representative images of mitochondria in young and aged cells. Images were acquired by electron microscopy (top) and by confocal microscopy, after staining of mitochondria with TOMM20 (bottom); (n=3 biological replicates for each age group). **(C)** Quantification of mtDNA content in young and aged cells by qPCR analysis of the expression levels of the mitochondrial-encoded a*tp6* and *cox-1* genes. Expression levels of the nucleus-encoded β-actin was used as an internal control, for normalization; (n=3 biological replicates for each age group). **(D)** Mean Fluorescence Intensity of young and aged cells after staining with MitoTracker Deep Red FM dye. Analysis was performed by flow cytometry; (n=3 biological replicates for young mice; n=2 biological replicates for old mice). **(E and F)** Schematic graph showing the two major sources of acetyl-CoA and the respective enzymes, which catalyze this conversion. Representative immunoblots for these two enzymes in young and aged MSCs. β-actin was used as a loading control. Data are presented as mean ± SEM. Statistical significance was determined using student’s t-Test; * p-value <0,05.

**Figure S4.**
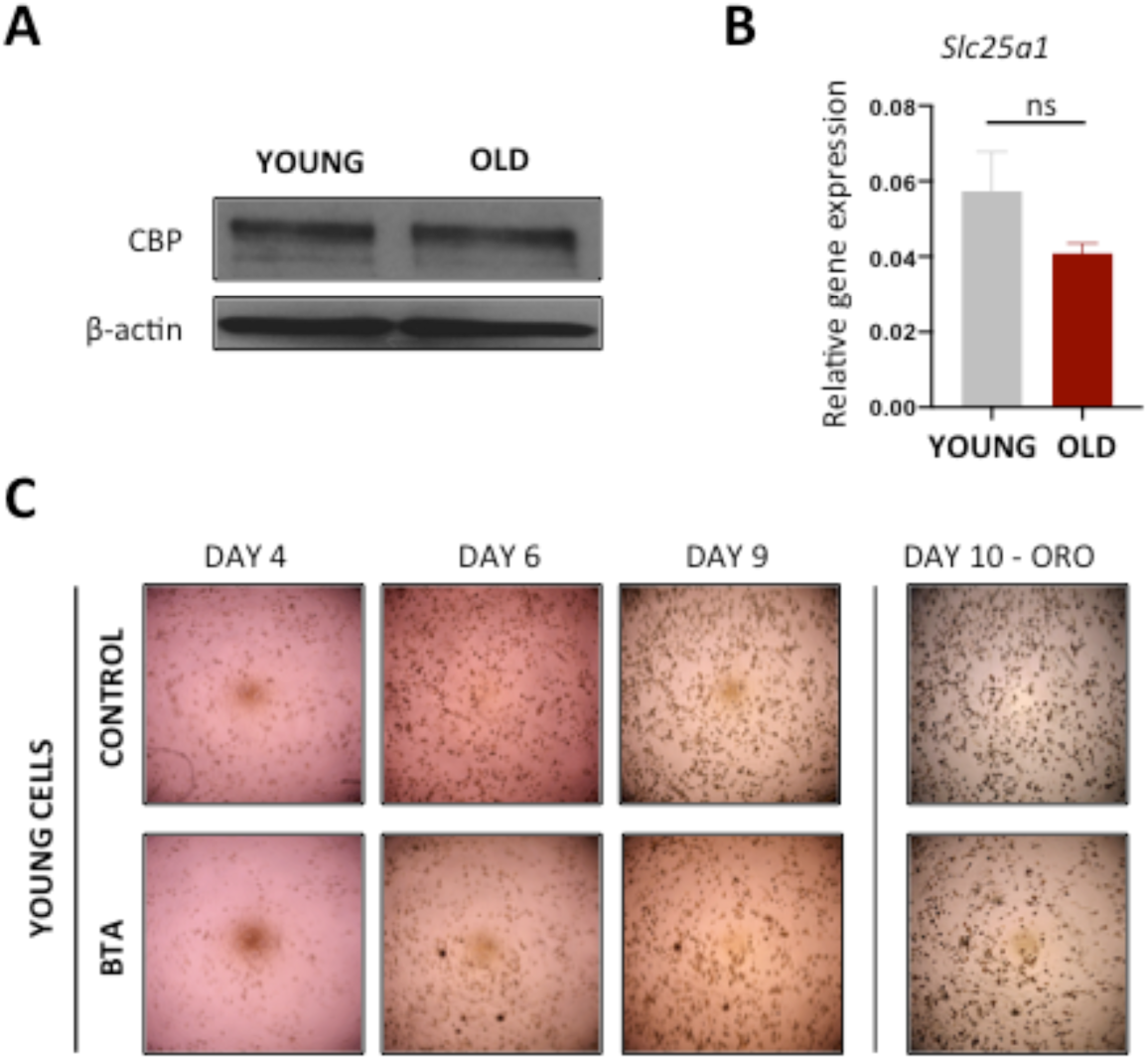
(related to Figure 4) **(A)** Representative immunoblots for the CBP histone acetyl-transferase in young and aged MSCs. β-actin was used as a loading control (n=2 for each age group). **(B)** qPCR analysis of the expression levels of the *Slc25a1* gene, which encodes the citrate carrier. β-actin expression levels were used as an internal control, for normalization; (n=3 biological replicates for young mice; n=2 biological replicates for old mice). **(C)** Representative images of adipocytes after 4 days, 6 days and 9 days of differentiation. Oil-Red-O staining performed on young MSCs, 10 days after induction of adipogenesis. Cells were treated with or without 1mM BTA during the adipogenic period (n=3 biological replicates for each group).

